# Interferon Dependent Immune Memory during HSV-1 Neuronal Latency Results in Increased H3K9me3 and Restriction of Reactivation by ATRX

**DOI:** 10.1101/2025.01.28.633013

**Authors:** Abigail L. Whitford, Gaelle Auguste, Alison K. Francois, Chris Boutell, Clint Miller, Sarah Dremel, Anna. R Cliffe

**Author notes:** Correspondence to Anna R. Cliffe.

## Abstract

Herpes simplex virus-1 (HSV-1) establishes a lifelong latent infection in neurons and reactivation from this latent state is the cause of recurrent oral and ocular infections, herpes simplex keratitis, and encephalitis. Neuronal conditions during initial HSV-1 infection have a long-term impact on latency, modulating how responsive latent genomes are to reactivation and, therefore, their ability to cause disease. Type I interferon (IFNα) exposure during initial infection results in promyelocytic leukemia nuclear-body (PML-NB) formation and a more restrictive form of HSV-1 latency. Here we demonstrate that IFNα induced PML-NBs recruit histone chaperones to the viral genome and promote the deposition of the repressive heterochromatin mark, histone H3 lysine 9 tri-methylation (H3K9me3), and its reader, ATRX (alpha-thalassemia/mental retardation, X-linked). This work reveals the mechanism by which immune signaling during initial infection induces an epigenetic memory on HSV-1 genomes that is maintained during latency and inhibits reactivation. ATRX is highly abundant in neurons and is essential for maintaining cellular heterochromatin during neuronal stress. Here we find that ATRX prevents transcription, and subsequent reactivation, from H3K9me3-bound latent genomes by remaining associated with viral chromatin following stress-induced phosphorylation of histone H3. This indicates that H3K9me3-associated viral genomes are refractory to reactivation when read by ATRX. This work demonstrates that ATRX acts as a neuronal restriction factor against HSV-1 reactivation, elucidating a new potential target for inhibiting HSV-1 reactivation and subsequent human disease.

## Introduction

Herpes simplex virus 1 (HSV-1) establishes a lifelong latent infection in neurons, allowing the virus to persist in its host for extended periods and evade immune surveillance. The DNA virus genome persists in the nucleus of an infected neuron and is subjected to heterochromatin-based silencing (Cliffe et al., 2009; Kwiatkowski et al., 2009; Nicoll et al., 2016; Wang et al., 2005). It is thought that epigenetic silencing mechanisms promote and maintain latency and need to be remodeled for reactivation to occur. However, studies investigating the contribution of heterochromatin to HSV-1 latency are complicated by the heterogeneous nature of the latent viral epigenome. For example, there is evidence of heterogeneity in the subnuclear distribution of viral genomes, levels of viral gene, expression, and reactivation (Catez et al., 2012; Dochnal et al., 2022; Ma et al., 2014; Maroui et al., 2016; Sawtell & Thompson, 2004). How different epigenetic structures form and whether they differentially contribute to reactivation competencies of the viral genome is unknown. This is important to understand in the context of HSV-1, as reactivation is associated with a variety of diseases, including cold sores, herpes simplex keratitis, and encephalitis. Further, studying the epigenetics of HSV infection of neurons also informs on the function of heterochromatin-associated proteins in this specialized cell type.

As a virus that persists in a long-lived cell type, the state of the cell at the time of initial infection could impact the nature of latency and future ability to reactivate. For example, neuronal stress or immune signaling during initial infection has been shown to modulate later reactivation (Dochnal et al., 2023; Suzich et al., 2021). The mechanisms underlying cell-intrinsic long-term memory during HSV-1 latency, triggered by short-term external stimuli, remain unknown. In the case of innate immune stimuli, our laboratory has previously shown that type I interferon exposure at the time of primary infection restricts later reactivation. We also showed that this was due to the formation of Promyelocytic leukemia-nuclear bodies (PML-NBs) in neurons (Suzich et al., 2021). PML-NBs are membrane-less nuclear organelles composed of the scaffolding protein PML and various stably and transiently associated proteins (Lallemand-Breitenbach & de The, 2010). PML-NBs function to compartmentalize proteins within eukaryotic cells and are known to play a role in the intrinsic repression of productive herpesvirus replication (Alandijany et al., 2018; Everett et al., 1998; Everett & Murray, 2005; Everett et al., 2004). A subpopulation of HSV genomes colocalizes with PML-NBs during latency (Catez et al., 2012). Our previous work showed that the co-localization of viral genomes with PML-NBs occurred when neurons were exposed to type I IFN solely at the time of initial infection and that PML was responsible for the IFN-dependent restriction of reactivation (Suzich et al., 2021). However, the mechanism underlying this restriction remains unclear, specifically whether it is due to changes in epigenetic silencing or the physical entrapment of viral genomes within PML-NBs.

The latent HSV-1 genome is known to be associated with repressive histone modifications. However, the intersection between extrinsic stimuli and the latent epigenome, both in the context of latency establishment and reactivation, are unknown. Latent viral genomes are enriched with H3K9me2, H3K9me3, and H3K27me3 (Cliffe et al., 2009; Kwiatkowski et al., 2009; Nicoll et al., 2016; Wang et al., 2005). H3K9me3 is viewed as a long-term, constitutive form of heterochromatin, whereas H3K27me3 is known to mark regions of facultative heterochromatin on the host, which can potentially more readily convert to active chromatin for gene expression (Dochnal et al., 2021). However, in the context of HSV-1 infection of a terminally differentiated neuron, it is unknown whether certain types of histone post-translational modifications give rise to genomes that are more or less capable of reactivating. Furthermore, investigating heterogeneity can be a challenge as most techniques analyze a population of genomes. To investigate heterochromatin marks on a single genome, we developed a tool (NucSpotA) to quantify the labeling of an immunofluorescent stain at the viral genome that can then be standardized to the stain throughout the nucleus. This tool has enabled us to quantify subpopulations of genomes that display distinct marks (Francois et al., 2023). We hypothesized that interferon-induced PML-NBs may increase the subpopulation of latent genomes with H3K9me3 silencing. This hypothesis is based on previous studies showing that PML-NBs impact H3K9me3 deposition on cellular chromatin (Delbarre et al., 2017) and potentially on viral DNA during lytic infection (Lee et al., 2016). It remains uncertain how an increase in the percentage of viral genomes with H3K9me3 would lead to a more repressive state of latency.

H3K9me3 represses chromatin through its interaction with histone readers, including HP1 family proteins (Bannister et al., 2001; Jacobs et al., 2001; Lachner et al., 2001), ATRX (alpha-thalassemia/mental retardation, X-linked) (Dhayalan et al., 2011; Noh et al., 2015), CHD4 (Mansfield et al., 2011), TRIM66 (Jain et al., 2020), PHRF1 (Jain et al., 2020), UHRF1 (Nady et al., 2011), TNRC18 (Zhao et al., 2023), MPP8 (Chang et al., 2011), and Tip60 (Sun et al., 2009), whereas the Polycomb family of protein complexes reads H3K27me3 (Cao et al., 2002). The diversity in reader proteins and the impact on chromatin structure increases the potential complexity of viral genome silencing. The readers associated with the latent HSV-1 genome remain unidentified, but understanding their role in silencing is crucial, as reader-based silencing would need to be overcome for reactivation to occur. HSV-1 reactivation is biphasic (Cliffe et al., 2015; Cuddy et al., 2020; Dochnal et al., 2022; Kim et al., 2012; Whitford et al., 2022) and the first phase of reactivation involves the phosphorylation of the serine 10 neighboring H3K9me3 (H3K9me3pS10). This phosphorylation evicts many H3K9me3 readers allowing for transcription without the removal of repressive methylation marks. Importantly, ATRX is an H3K9me3 reader that has been shown previously to be resistant to eviction following S10-phosphorylation (Noh et al., 2015). We investigated whether ATRX restricts HSV-1 reactivation and found that ATRX inhibits reactivation from H3K9me3-enriched genomes. This inhibitory effect is further amplified by IFN treatment, which increases ATRX enrichment at the latent viral genome.

## Results

### PML is required for the initiation but not the maintenance of the type I IFN-dependent restricted form of HSV-1 latency

Previously, using an *in vitro* model system of HSV-1 latent infection, we reported that PML-NBs form in peripheral sympathetic and sensory neurons following exposure to type I IFN. We also observed that the HSV-1 genome co-localizes and is maintained at PML-NBs only when initial infection occurs at the same time as exposure to type I IFN (Suzich et al., 2021). Type I IFN treatment during initial infection also resulted in repression of reactivation that could be overcome by the depletion of PML before infection. To determine the mechanism of PML-based restriction, we asked whether PML was required for both the initiation and maintenance of the repressive reactivation phenotype. Primary neurons from the superior cervical ganglia (SCG) of postnatal mice were dissected and transduced with lentivirus expressing *Pml* shRNA five days before infection with HSV-1. Infection was carried out using Stayput GFP (MOI 5 PFU/cell) (Dochnal et al., 2022), which expresses a Us11-GFP and is also defective in cell-to-cell spread, permitting the quantification of individual reactivating neurons (Figure 1A). Neurons were also treated with 600 IU/μl of IFNα for 18h prior to infection. Acyclovir (ACV) was added for the first 8 days post-infection to promote latency establishment and then washed out. Reactivation was triggered 10 days post-infection using the PI3K inhibitor, LY294002, as described previously (Camarena et al., 2010; Cliffe et al., 2015; Dochnal et al., 2022; Kim et al., 2012; Kobayashi et al., 2012) (Figure 1A). Consistent with our previous study (Suzich et al., 2021), we found PML depletion prior to infection resulted in enhanced reactivation of IFN-treated neurons to levels comparable to the untreated neurons (Figure 1B). However, depletion of PML after latency had already been established (5 days post-infection) did not impact reactivation in IFN-treated neurons (Figure 1B). These findings suggest that the presence of PML-NBs during initial infection contributes to the restrictive phenotype initiated by IFNα treatment during infection, but PML-NBs were not required to maintain IFNα-induced silencing at later time points.

**Figure 1:**
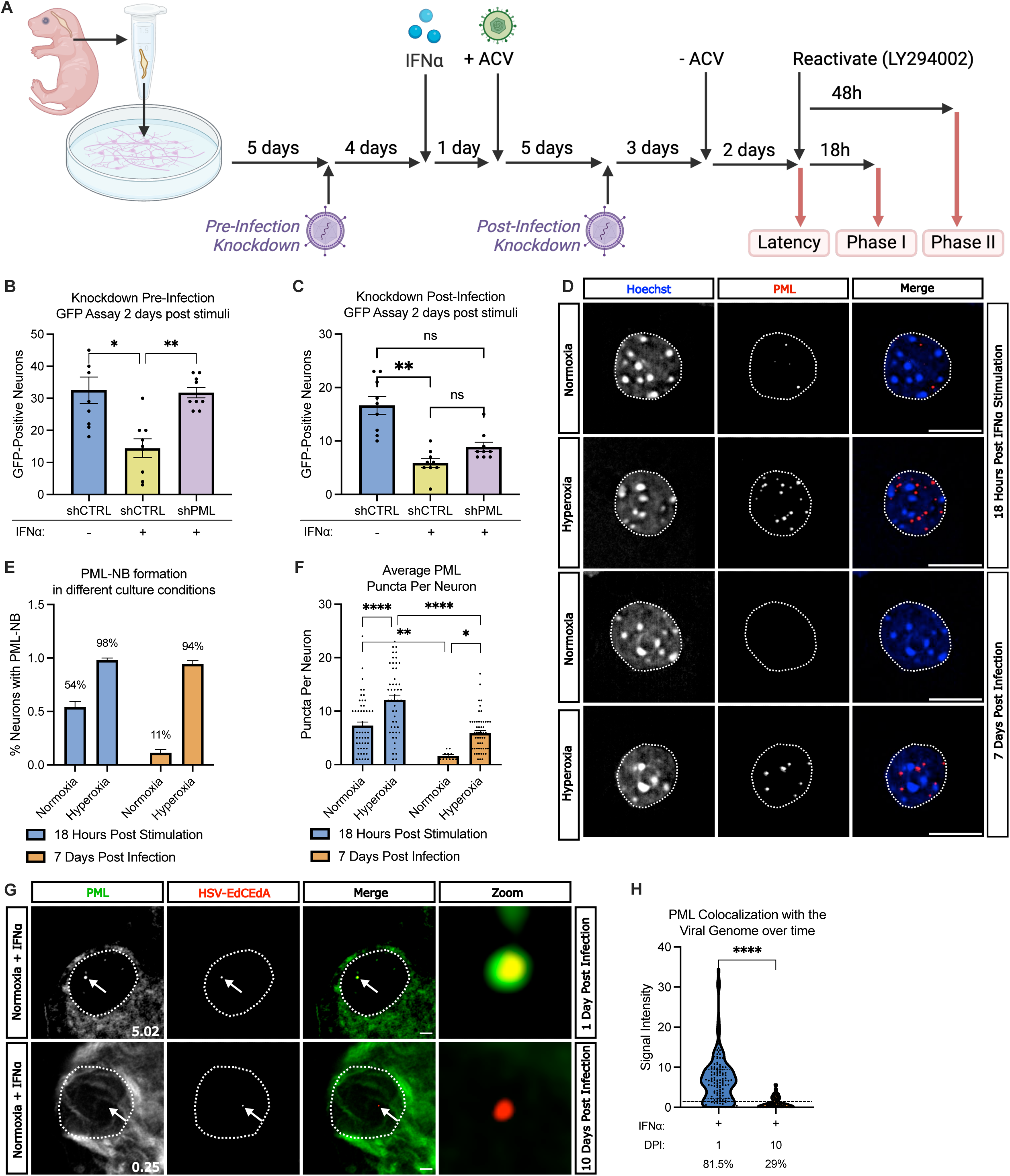
PML-NBs initiate neuronal HSV-1 silencing but do not persist to latency to maintain silencing. **A)** Schematic of the primary neuronal model of HSV-1 latency and reactivation using sympathetic neurons isolated from the superior cervical ganglia (SCG) of newborn mice. **B)** Quantification of HSV-1 reactivation based on the numbers of Us11-GFP positive neurons at 2 days post reactivation. Neurons were depleted of PML using lentivirus-mediated shRNA depletion five days prior to infection and treated with 600 IU/ml IFNα for 18h hours before infection and 24 hours post-infection. Reactivation was induced using LY294002 (20 μM). Individual repetitions (N=9; wells containing approximately 5000 neurons) from 3 independent dissections. Statistical comparisons were made using an Ordinary one-way ANOVA with Tukey’s multiple comparison. **C)** Quantification of HSV-1 reactivation based on the numbers of Us11-GFP positive neurons at 2 days post reactivation. Neurons were treated IFNα before infection as in B and depleted of PML using lentivirus-mediated shRNA depletion five days post-infection. N=9 biological replicates from 3 independent dissections. Statistical comparisons were made using an Ordinary one-way ANOVA with Tukey’s multiple comparison. **D)** Representative images of sympathetic neurons cultured in atmospheric (21% Oxygen) or physioxia (5% oxygen) conditions and fixed at either 18 hours post-treatment with IFNα (or 0h post-infection) or 7 days post-infection. Immunofluorescence was carried out for PML and cells co-stained with Hoechst. Scale bar, 10 μm. **E & F)** Quantification of the percentage of neurons with detectable PML-NBs in their nuclei (E) and (F) detectable PML puncta per nuclei at 18 hours post-treatment with IFNα or 7 days post-infection in either atmospheric or physioxia conditions. N>50 cells from 2 biological replicates. Statistical comparisons were made using a two-way ANOVA with Tukey’s multiple comparisons. In F, dots represent individual cells. **G)** Neurons treated with IFNα at the time of infection were infected with HSV-1 EdC/EdA and fixed at 1- or 10-days post-infection. The viral genome was visualized using click chemistry, and immunofluorescence was carried out for PML. White arrows point to location of viral genome. Scale bar, 10 μm. **H)** NucSpotA was used to quantify the signal intensity of PML at the viral genome 1- or 10-days post-infection. Each data point represents one viral genome. Percentages indicate the proportion of genomes with NucSpotA intensity ratios above the denoted co-localization threshold (dashed line). N>60 cells from 3 biological replicates. Statistical comparisons were made using a Mann-Whitney. Data represent the mean ± SEM. (ns not significant, *≤ 0.05, **≤ 0.01, ***≤ 0.001, ****≤ 0.0001).

The contribution of PML to the maintenance of the restrictive phenotype differed slightly from our previous study, where we had observed a slight increase in reactivation in neurons following PML depletion after latency was established. In our original study, neurons were cultured in atmospheric conditions. However, we have since changed the culturing conditions to more accurately mimic the oxygen concentration of peripheral neurons, which improves neuronal health, particularly in long-term cultures (Dochnal et al., 2024). We investigated whether the change in culturing conditions impacted the formation and maintenance of PML-NBs. We found that IFNα treated neurons cultured in more physiological oxygen levels (5% O_2_) developed fewer PML-NBs and had decreased PML-NB persistence than those cultured in atmospheric oxygen (Figure 1F). These data suggest that the oxygen concentration that neurons are cultured in *in vitro* affects their response to innate immune signals. Moreover, although PML initially co-localized with HSV-1 following IFNα treatment in both atmospheric and physiological oxygen concentrations, after 10 days the co-localization was lost under physioxia. We quantified this phenotype over multiple latent neurons using NucSpotA (Francois et al., 2023) (Figure 1H) and observed 81.5% of genomes co-localized at 1 day in contrast to 29% at 10 days. The percentage of genomes co-localizing was determined by setting a signal intensity threshold based on colocalization by eye on 50 genomes. Together, our data indicate that PML associates with the viral genome during initial infection and initiates silencing, but PML-NBs do not persist during latency and are not required to maintain the restrictive phenotype of IFNα.

### IFNα induced restriction during initial infection is DAXX dependent

Our observation that PML was not required to maintain the silent state induced by IFN treatment argued against a role for the PML protein in directly contributing to genome inaccessibility in latently infected neurons. Instead, this observation argued for a PML-dependent downstream impact on latent viral genomes. We hypothesized this was due to PML-dependent altered epigenetic structures resulting from type I IFN treatment. Therefore, we focused on PML-NB components that have known roles in heterochromatin targeting. We first investigated the contribution of Death Domain Associated Protein (DAXX) to the repressive phenotype because it is a constitutive component of PML-NBs and plays a role in the deposition of histones at regions of heterochromatin. Further, previous studies have found that DAXX represses HSV-1 lytic infection (Alandijany et al., 2018; Cabral et al., 2018; Lukashchuk & Everett, 2010) via its role as an H3.3 chaperone (Cabral et al., 2021; Cohen et al., 2018; Drane et al., 2010; Lewis et al., 2010), although the function of DAXX during HSV latent infection is unknown. Using two independent shRNAs, DAXX knockdown (Supplemental Figure 1) was able to recover reactivation in IFNα pre-treated neurons when it was knocked down before infection (Figure 2A & 2B). Consistent with the data on PML, the viral genome also co-localized with DAXX at early time points (1 day) following infection in neurons treated with type I IFN (Figure 2C) and the co-localization of viral genomes with DAXX was not maintained during latent infection arguing against a role for DAXX in maintaining the repressive nature of HSV-1 latency (Figure 2D & 2E). Additionally, we sought to determine whether the presence of PML was required for the observed enrichment of DAXX. We found that depletion of PML before infection significantly reduced the association of viral genomes with DAXX (from 80.5% of genomes co-localized with DAXX to 0.18%) (Figure 2F), demonstrating DAXX contributes to the repressive effects of IFNα and is recruited to viral genomes in a manner that is dependent on PML.

**Figure 2:**
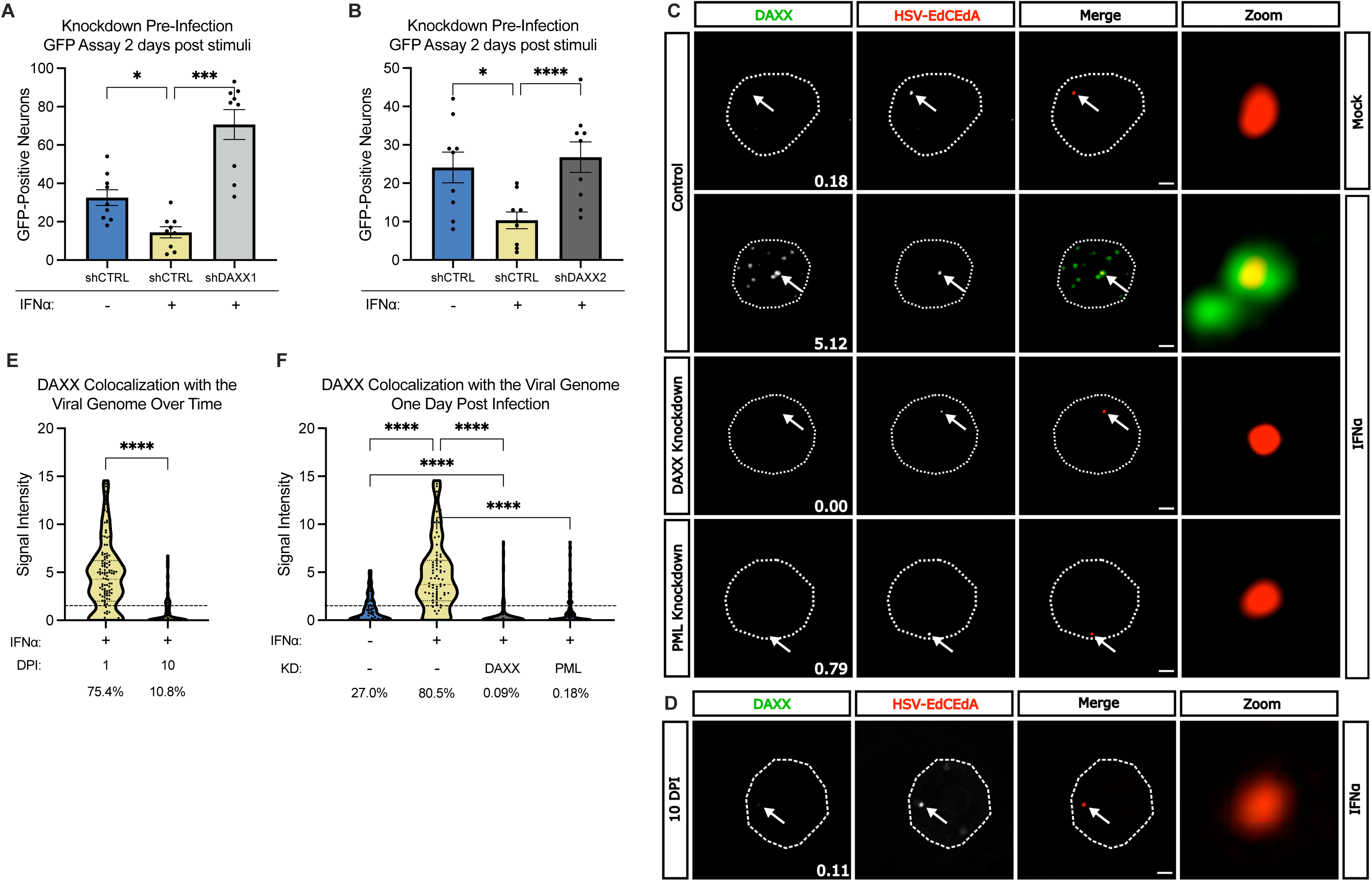
IFNα induced restriction during initial infection is DAXX dependent. **A & B**) Sympathetic neurons were infected with Stayput HSV-1 in the presence or absence of IFNα and depleted of DAXX using shRNAs at 5 days pre-infection using two independent shRNAs. Reactivation was quantified based on the numbers Us11-GFP expressing neurons following addition LY294002. Data represent the mean ± SEM. N=9 biological replicates from 3 independent dissections. Statistical comparisons were made using an Ordinary one-way ANOVA with Tukey’s multiple comparison. **C & D)** Representative images of sympathetic neurons untreated or treated with 600 IU/ml of IFNα cultured in 5% oxygen. Neurons were infected with EdC/EdA labeled HSV-1 and depleted of PML or DAXX at 5 days pre-infection. Cells were fixed at 1 day (C) and 10 days (D) post-infection. The viral genome was visualized using click chemistry, and immunofluorescence was carried out for DAXX. White arrows point to location of viral genome. Scale bar, 10 μm. **E & F)** NucSpotA was used to quantify the signal intensity of DAXX at the viral genome 1- or 10-days post infection. Each data point represents one viral genome. Percentages indicate the proportion of genomes with NucSpotA intensity ratios above the denoted co-localization threshold (dashed line). N>60 cells from 3 biological replicates. Statistical comparisons were made using **E)** Mann-Whitney and **F)** Kruskal-Wallis test with Dunn’s multiple comparison. (ns not significant, *≤ 0.05, **≤ 0.01, ***≤ 0.001, ****≤ 0.0001)

### IFNα stimulation increases H3K9me3 enrichment on the viral genome that is maintained during latency

Previous studies have demonstrated that PML-NBs contribute to increased enrichment of H3K9me3 on cellular chromatin (Delbarre et al., 2017) and potentially on HSV-1 during lytic infection (Francois et al., 2023). Additionally, DAXX has been associated with the formation of heterochromatin characterized by H3K9me3 (Voon et al., 2015), which is known to be enriched on populations of HSV-1 genomes during quiescent infection (Cliffe et al., 2009) (Cohen et al., 2018; Roubille et al., 2024). We hypothesized that type I IFN exposure alters the epigenetic silencing of the latent HSV-1 genome via increased H3K9me3 in a DAXX and PML-dependent manner.

To test this hypothesis, we carried out immunofluorescence to determine the colocalization of H3K9me3 with the viral genome once latency was established (Figure 3A). To overcome problems in quantifying the co-localization of nuclear proteins with viral genomes and take into account the heterogeneity of histone enrichment at different viral genomes in a population, we used NuncSpotA; a program that quantifies the labeling of an immunofluorescent signal at the viral genome and standardizes that enrichment with the stain throughout the nucleus (Francois et al., 2023). Using this approach, we identified a subpopulation of viral genomes that co-localized with H3K9me3 in untreated neurons (49.3%), which increased in the IFNα treatment to 75.3% (Figure 3B). Depletion of either PML or DAXX prior to infection reduced the proportion of viral genomes co-localized with H3K9me3 to levels similar to untreated neurons (30.2% for PML depletion and 33.3% for DAXX depletion). We did not observe a significant decrease in H3K9me3 enrichment when PML or DAXX depletion occurred after latency had been established (55.9% in PML-depleted and 51.5% in DAXX-depleted neurons). These data indicate that viral genomes co-localize with H3K9me3 during latent infection, and this co-localization increases when type I IFN is present during initial HSV-1 infection. Further, in the presence of type I IFN, the increased H3K9me3 co-localization was both PML and DAXX-dependent during early infection but were not required for the mark to persist at later time points.

**Figure 3:**
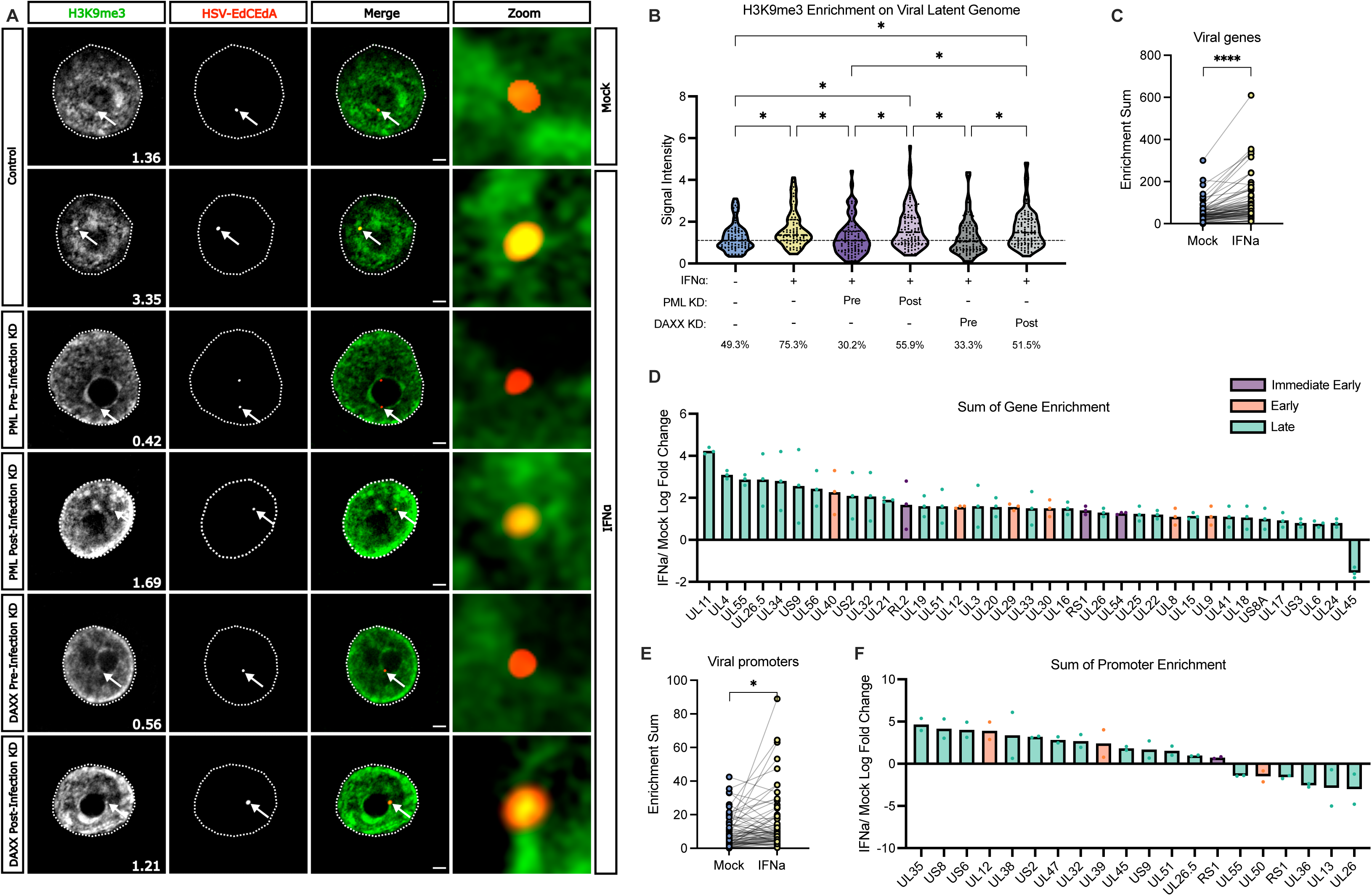
IFNα stimulation increases H3K9me3 enrichment on the viral genome that is maintained during latency. **A)** Representative images of sympathetic neurons untreated or treated with 600 IU/ml of IFNα cultured in 5% oxygen. Neurons were infected with EdC/EdA labeled HSV-1 and depleted of PML or DAXX at 5 days pre-infection or 5 days post-infection. Cells were fixed at 10 days post-infection. The viral genome was visualized using click chemistry, and immunofluorescence was carried out for H3K9me3. White arrows point to location of viral genome. Scale bar, 10 μm. **B)** NucSpotA was used to quantify the signal intensity of H3K9me3 at the viral genome 10-days post-infection. Each data point represents one viral genome. Percentages indicate the proportion of genomes with NucSpotA intensity ratios above the denoted co-localization threshold (dashed line). N>60 cells from 3 biological replicates. Statistical comparisons were made using a repeated Measure One way ANOVA. **C)** Sympathetic neurons were infected Stayput-GFP HSV-1 at an MOI of 10 PFU/cell in the presence or absence of IFNα. CUT&Tag was performed at 10 days post-infection using an antibody against H3K9me3. Fragments were sequenced and aligned to the viral genome. The sum of coverage at the defined gene body and promoter regions were used to calculate the enrichment of H3K9me3. The average enrichment sum of all viral gene bodies is plotted, with a line connecting the mock and IFNα treatments for individual viral open reading frames. Statistical analysis was made using a two-way ANOVA. N=2 biological replicates **D)** Plot of gene bodies with a log fold change of greater than 0.5 or less than -0.5 across both replicates. Immediate early genes are colored purple, early genes are colored orange, and late genes are colored teal. **E)** The average enrichment sums of all viral gene promoters. The line connects the same viral promoter in mock and IFNα treated cultures. Statistical analysis was made using a two-way ANOVA. **F)** Plot of gene promoters that had a log fold change of greater than 0.5 or less than -0.5 across both replicates. Immediate early genes are colored purple, early genes are colored orange, and late genes are colored teal. **G)** Representative integrative genome viewer images. All data was normalized to total mapped reads. Viral genes are shown in blue and viral promoters are shown in red. The y-axes were group auto-scaled between conditions. (ns not significant, *≤ 0.05, **≤ 0.01, ***≤ 0.001, ****≤ 0.0001).

To further validate the increased H3K9me3 association upon IFNα treatment and investigate where on viral genomes H3K9me3 was enriched, we performed Cleavage Under Targets and Tagmentation (CUT&Tag) for H3K9me3 on latently infected neurons. Consistent with our immunofluorescence results, IFNα treatment significantly increased H3K9me3 enrichment over viral gene bodies (Fig. 3C, 3D) as well as at promoters (Fig. 3E, 3F). Viral gene bodies and promoters were quantified as H3K9me3 enrichment at both cellular promoters and cellular gene bodies has been correlated with epigenetic repression and transcriptional silencing (Lee et al., 2020). The sum of H3K9me3 enrichment over viral gene bodies (excluding promoter regions) and over promoters was quantified. Genes that had at least a twofold increase in H3K9me3 over two replicates were plotted. We concluded that there were consistent regions of the genome with increased H3K9me3 in IFNα treated. These findings suggest that IFNα promotes more H3K9me3 on the genome and that this mark persists during latency and correlates with a decrease in reactivation.

### ATRX inhibits HSV-1 reactivation, and this inhibition is enhanced by IFNα treatment

Given our observation that PML and DAXX contributed to the initiation of IFN-mediated repression via H3K9me3, we set out to investigate ATRX, a known interaction partner of DAXX. ATRX complexes with DAXX to deposit H3.3 and, via its interaction with DAXX, represses lytic HSV-1 infection in human fibroblasts (Lukashchuk & Everett, 2010) (Cabral et al., 2021). ATRX can also function as an H3K9me3 reader (Dhayalan et al., 2011) and has been shown to maintain the enrichment of H3K9me3 on lytically replicating viral genomes (Cabral et al., 2018). However, the role of ATRX in HSV-1 gene silencing during latency is unknown. We depleted ATRX prior to HSV infection and once latency had been established using two independent shRNAs (Supplemental Figure 4A, 4B, & 4C). We found ATRX depletion prior to infection resulted in enhanced reactivation of IFN-treated neurons (Figure 4A & 4B). In contrast to PML and DAXX, the depletion of ATRX after latency was established (Figure 4C & 4D), also resulted in significantly enhanced reactivation in IFN-treated neurons. This finding indicates that unlike PML and DAXX, ATRX is required for maintaining the restrictive phenotype in IFNα treated neurons.

**Figure 4:**
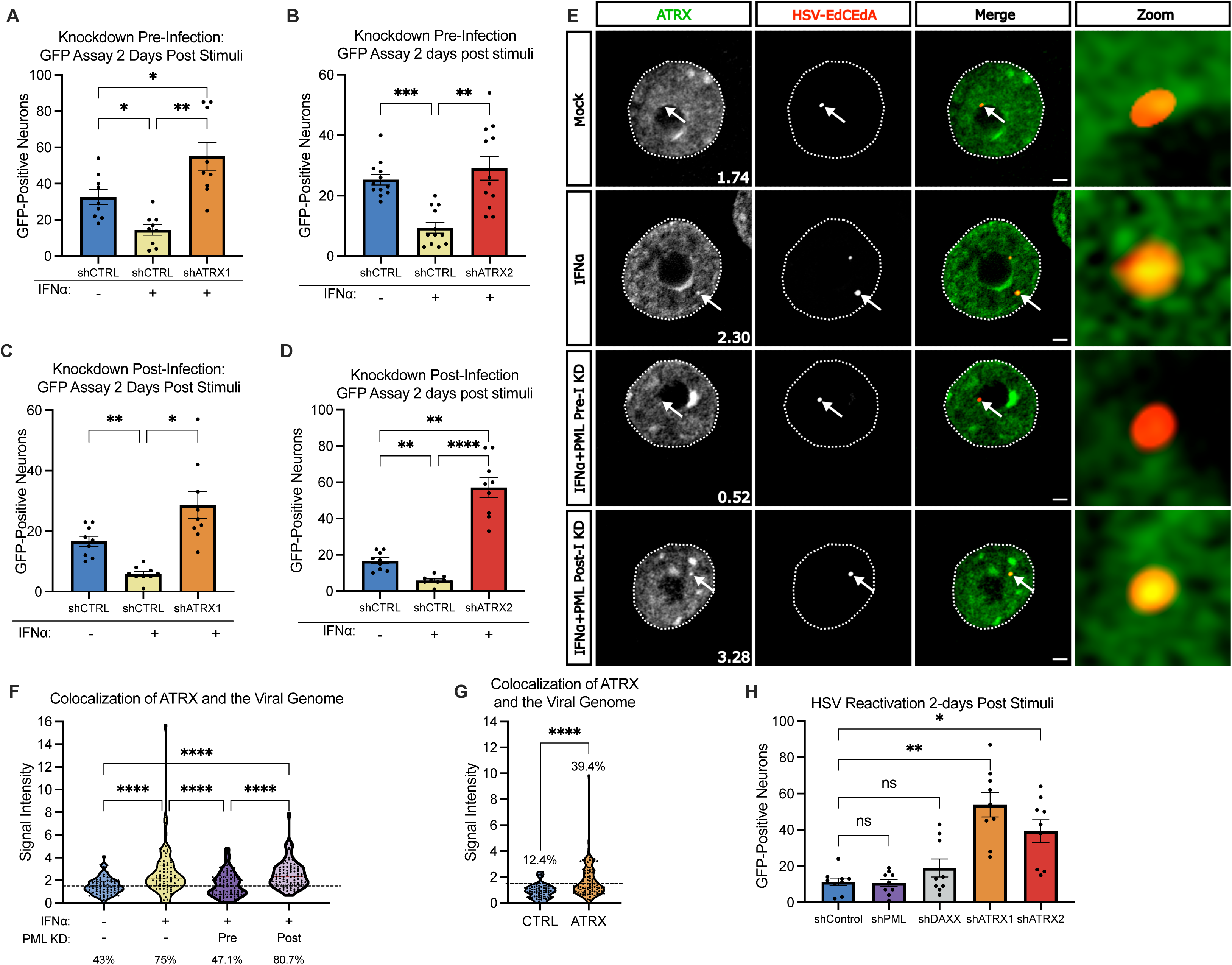
ATRX inhibits HSV-1 reactivation, and this inhibition is enhanced by IFNα treatment. **A & B**) Sympathetic neurons were infected with Stayput HSV-1 in the presence or absence of IFNα and depleted of ATRX using shRNAs at 5 days pre-infection using two independent shRNAs. Reactivation was quantified based on the numbers Us11-GFP expressing neurons following addition LY294002. Data represent the mean ± SEM. N=9 biological replicates from 3 independent dissections. Statistical comparisons were made using an Ordinary one-way ANOVA with Tukey’s multiple comparison. **C & D**) Sympathetic neurons were infected with Stayput HSV-1 in the presence or absence of IFNα and depleted of ATRX using shRNAs at 5 days post-infection using two independent shRNAs. Reactivation was quantified based on the numbers Us11-GFP expressing neurons following addition LY294002. Data represent the mean ± SEM. N=9 biological replicates from 3 independent dissections. Statistical comparisons were made using an Ordinary one-way ANOVA with Tukey’s multiple comparison. **E)** Representative images of sympathetic neurons untreated or treated with 600 IU/ml of IFNα cultured in 5% oxygen. Neurons were infected with EdC/EdA labeled HSV-1 and depleted of PML at 5 days pre-infection or 5 days post-infection. Cells were fixed at 10 days post-infection. The viral genome was visualized using click chemistry, and immunofluorescence was carried out for ATRX. White arrows point to location of viral genome. Scale bar, 10 μm. **F)** NucSpotA was used to quantify the signal intensity of ATRX at the viral genome 10 days post-infection. Each data point represents one viral genome. Percentages indicate the proportion of genomes with NucSpotA intensity ratios above the denoted co-localization threshold (dashed line). N>60 cells from 3 biological replicates. Statistical comparisons were made using Kruskal-Wallis test with Dunn’s multiple comparison. **G)** NucSpotA was used to quantify the signal intensity of ATRX at the viral genome 10 days post-infection in mock-treated neurons. CTRL denotes a randomized control where the genome-containing layer is rotated 180 degrees to provide a value for random placement. Each data point represents one viral genome. Percentages indicate the proportion of genomes with NucSpotA intensity ratios above the denoted co-localization threshold (dashed line). N>60 cells from 3 biological replicates. Statistical comparisons were made using an unpaired t-test. **H)** Sympathetic neurons were infected with Stayput HSV-1 in the absence of IFNα and depleted of PML, DAXX, or ATRX using shRNAs at 5 days post-infection. Reactivation was quantified based on the numbers Us11-GFP expressing neurons following addition LY294002. Data represent the mean ± SEM. N=9 biological replicates from 3 independent dissections. Statistical comparisons were made using an Ordinary one-way ANOVA with Tukey’s multiple comparison. (ns not significant, *≤ 0.05, **≤ 0.01, ***≤ 0.001, ****≤ 0.0001).

Supporting a role for ATRX in maintaining the repressive nature of HSV-1 latency, we found that unlike DAXX and PML, ATRX co-localization with the viral genome showed detectible co-localization with ATRX at 10 days post-infection, indicating that ATRX is maintained on viral genomes once latency has been established (Figure 4E). This colocalization significantly increased with IFNα treatment (from 43% of genomes co-localized with ATRX to 75%) (Figure 4F). Notably, this association level was similar to the proportion also associated with H3K9me3, as shown in Figure 3B (49.3% in untreated and 75.3% type I IFN treated). We then sought to determine whether the presence of PML was required for the observed increase in the enrichment of ATRX. We found that depletion of PML before infection significantly reduced the association of viral genomes with ATRX (from 75% of genomes co-localized to 47.1%) in IFN-pretreated neurons. However, depletion of PML after infection did not impact ATRX enrichment. Therefore, the IFN-dependent increase in ATRX association with the viral genome was dependent on PML during initial infection but did not require PML for the maintained association. These data further support that IFNα induced PML-NBs enact long-lasting epigenetic changes that are maintained independently of the bodies themselves during latency.

In contrast to PML and DAXX, we found that ATRX co-localized viral genomes even in the untreated neurons (43%), albeit to lower levels than IFN-treated neurons (Figure 4F). To determine whether this association occurred at levels above those obtained for random co-localization, we compared the co-localization values to those of a rotated control and found that ATRX was significantly enriched on the viral genome (Figure 4G). This prompted us to investigate the role of ATRX in neurons that had not been treated with IFNα. ATRX, unlike PML and DAXX, is highly abundant in neurons of the central nervous system (Berube et al., 2005). We investigated the relative levels of ATRX in murine (SCG) and human (HD10.6 (Raymon et al., 1999)) peripheral neurons in contrast to murine (primary dermal fibroblasts) and human (human foreskin fibroblasts) fibroblasts (Supplemental Figure 4D). We chose to quantify by measuring immunofluorescence staining because of the challenges of normalizing cell numbers between vastly distinct cell types and problems carrying out Western blotting for ATRX in neurons where cell numbers are low. Consistent with previous studies, we observed higher staining intensity of ATRX in neuronal cell nuclei compared to non-neuronal cells (Supplemental Figure 4E). These results indicate that ATRX is highly abundant in neurons of the peripheral nervous system even without IFNα stimulation.

We investigated the impact of ATRX in repressing reactivation in untreated neurons. When ATRX was depleted after latency was established, we observed a significant increase in HSV-1 reactivation. We did not observe the same increase in reactivation following the depletion of PML or DAXX (Figure 4H), again consistent with ATRX playing a role in restricting HSV-1 reactivation in a manner independent of both PML and DAXX. We also observed a slight increase in the numbers of GFP-positive neurons following ATRX depletion alone in the absence of the addition of the reactivation trigger (Supplemental Figure 4F). This increase in GFP-positive neurons was very slight (from an average of 1.7 to 4) and was lower than the numbers observed with a reactivation trigger. We also found the increase in GFP could be blocked by an inhibitor of the DLK-dependent neuronal cell stress pathway. We hypothesized that the increase in GFP reflects an increase in spontaneous reactivation as a result of low levels of stress in the neuronal cultures as opposed to ATRX depletion directly inducing reactivation. In summary, our data indicate ATRX plays a role in limiting HSV reactivation, and its accumulation on viral genomes increases in IFN-pretreated neurons in a manner reliant on PML-NB formation.

### ATRX remains co-enriched with H3K9me3S10p at the viral genome during reactivation, preventing Phase 1 and protecting H3K9me3 bound genomes from undergoing full reactivation

Because we found ATRX functioned to inhibit HSV-1 reactivation in both mock and IFNα-treated conditions, and that ATRX enrichment levels mirrored those for H3K9me3, we hypothesized that ATRX restricted HSV-1 reactivation through its ability to read H3K9me3. To test this hypothesis, we performed immunofluorescence experiments to directly assess the co-enrichment of ATRX and H3K9me3 on the viral genome during latency at the single genome level (Figure 5A). By using NucSpotA with the previously defined cut-offs for positive co-localization, we could determine the co-enrichment values for H3K9me3 and ATRX. We found that treatment with interferon at the time of infection increased the population of neurons co-enriched with ATRX and H3K9me3 from 36% in untreated to 61% in IFNα pulsed (Figure 5B). Furthermore, in both mock and interferon conditions, we found only a small population of neurons with only H3K9me3 (14% in both conditions) or only ATRX (12% in mock and 16% in interferon pulsed). We determined that individual genomes that did not reach the co-localization threshold for one mark also did not reach the threshold for the second mark. Meanwhile, genomes positively co-localized with one tend to be co-localized with both. This pattern implies that the presence or absence of one factor may influence or depend on the other, reflecting interdependence or cooperative roles for ATRX and H3K9me3. These data imply ATRX functions as an H3K9me3 reader on a subset of viral genomes and with interferon treatment this subset increases.

**Figure 5:**
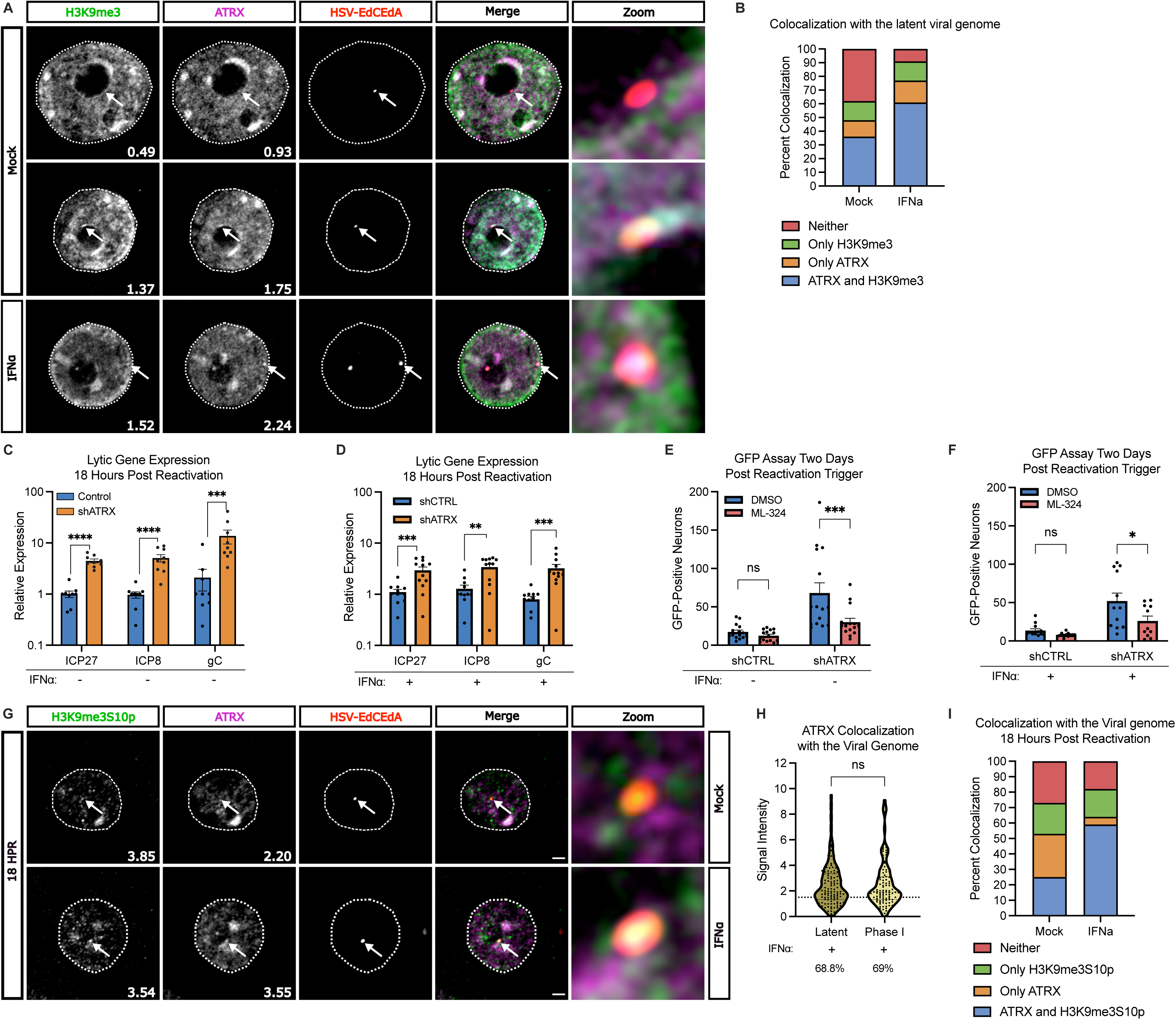
ATRX remains co-enriched H3K9me3S10p at the viral genome during reactivation, preventing Phase 1 and protecting H3K9me3 bound genomes from undergoing full reactivation. **A)** Representative images of sympathetic neurons untreated or treated with 600 IU/ml of IFNα cultured in 5% oxygen. Neurons were infected with EdC/EdA labeled HSV-1. Cells were fixed at 10 days post-infection. The viral genome was visualized using click chemistry, and immunofluorescence was carried out for H3K9me3 and ATRX. White arrows point to location of viral genome. Scale bar, 10 μm. **B)** NucSpotA was used to quantify the signal intensity of H3K9me3 and ATRX at the viral genome 10 days post-infection. Percentage of genomes above or below the denoted co-localization threshold (H3K9me3=1.1 and ATRX=1.5) are represented as a bar graph. N>60 cells from 3 biological replicates. **C & D**) Sympathetic neurons were infected with Stayput HSV-1 in the absence (C) or presence (D) of IFNα and depleted of ATRX using shRNAs at 5 days post-infection. RNA was collected 18 hours following the addition LY294002 (Phase I). RT-qPCR was used to quantify IE (ICP27), E (ICP8), and L (gC) viral gene expression. Statistical comparisons were made with a t test or Mann-Whitney test, Error bars: SEM) N=9 biological replicates from 3 independent dissections. **E & F**) Sympathetic neurons were infected with Stayput HSV-1 in the absence (E) or presence (F) of IFNα and depleted of ATRX using shRNAs at 5 days post-infection. Reactivation was quantified based on the numbers Us11-GFP expressing neurons following addition LY294002 and DMSO or ML-324 (10μM; JMJD2 inhibitor). Data represent the mean ± SEM. N=9 biological replicates from 3 independent dissections. Statistical comparisons were made using a two-way ANOVA with Šídák’s multiple comparisons test. **G)** Representative images of sympathetic neurons untreated or treated with 600 IU/ml of IFNα cultured in 5% oxygen. Neurons were infected with EdC/EdA labeled HSV-1. Cells were fixed 18 hours post-reactivation stimulus. The viral genome was visualized using click chemistry, and immunofluorescence was carried out for H3K9me3S10p and ATRX. White arrows point to location of viral genome. Scale bar, 10 μm. **H)** NucSpotA was used to quantify the signal intensity of ATRX at the viral genome 10 days post-infection or 18 hours post-reactivation stimulus. Each data point represents one viral genome. Percentages indicate the proportion of genomes with NucSpotA intensity ratios above the denoted co-localization threshold (dashed line). N>60 cells from 3 biological replicates. Statistical comparisons were made using a Mann Whitney. **I)** NucSpotA was used to quantify the signal intensity of H3K9me3S10p and ATRX at the viral genome 18 hours post-reactivation stimulus. The percentage of genomes above or below the denoted co-localization threshold (H3K9me3S10p=1.5 and ATRX=1.5) are represented as a bar graph. N>60 cells from 3 biological replicates. (ns not significant, *≤ 0.05, **≤ 0.01, ***≤ 0.001, ****≤ 0.0001)

We then investigated the enrichment of ATRX at viral genomes following a reactivation stimulus. Previous studies from our lab and others (Cliffe et al., 2015; Dochnal et al., 2022; Kim et al., 2012; Whitford et al., 2022) have established that reactivation of HSV-1 occurs in two distinct phases. Phase I is initiated when neurons are exposed to specific stressors, such as the loss of nerve growth factor signaling. During Phase I, there is JNK-dependent phosphorylation of the serine residue adjacent to lysine 9 on histone H3 (H3K9me3S10ph) (Cliffe et al., 2015; Cuddy et al., 2020). This "methyl/phospho switch" occurs on cellular chromatin during stress resulting in the eviction of some H3K9me3 readers that are unable to remain bound after serine 10 phosphorylation, allowing for transcription without the removal of post-translational modifications. However, ATRX exhibits the capability to bind to H3K9me3 even in the presence of serine 10 phosphorylation, maintaining cellular heterochromatic silencing (Noh et al., 2015). Although we have quantified reactivation following ATRX depletion based on GFP-positive neurons, this is a measure of full, Phase II reactivation. We first validated that ATRX specifically prevents entry into Phase I instead of preventing the progression from Phase I to Phase II. Depletion of ATRX after latency was established resulted in increased viral lytic gene expression in all viral gene classes (ICP27, ICP8, and gC) at the Phase I time point (18 hours post-reactivation). This ATRX-dependent increase was observed in both mock-treated (Figure 5C) and IFNα pre-treated neurons (Figure 5D) and was independent of DAXX as depletion of DAXX after latency had been established did not impact Phase I gene expression (Supplemental Figure 5A). Therefore, ATRX acts independently of DAXX to specifically prevent the exit of latent genomes into Phase I gene expression following a reactivation stimulus.

Phase I of reactivation is followed by Phase II, which involves the removal of repressive PTMs, replication of viral DNA, and virus production. The removal of histone PTMs H3K27me3 and H3K9me2 is required for Phase II of reactivation (Cliffe et al., 2015; Dochnal et al., 2022; Kim et al., 2012; Whitford et al., 2022). However, using a H3K9me3 demethylase (KDM4) inhibitor (ML-324), we found that the removal of H3K9me3 does not contribute to reactivation in both mock (Figure 5E) and IFNα (Figure 5F) treated neurons. Interestingly, only when ATRX is depleted before reactivation does inhibiting H3K9me3 removal contribute to reactivation. This further supports the hypothesis that ATRX functions as a reader of H3K9me3 on latent genomes, preventing H3K9me3-associated genomes from reactivating.

To investigate the mechanism by which ATRX prevents H3K9me3-associated genomes from reactivation, we asked whether H3K9me3 enriched genomes were still phosphorylated during Phase I when associated with ATRX. We performed immunofluorescence on HSV-1 infected neurons, either untreated or treated with IFNα during initial infection. (Figure 5G). ATRX enrichment on the viral genome was not significantly different at latent and Phase I time points (Fig 5D). Furthermore, quantification of both ATRX and H3K9me3pS10 at viral genomes using NucSpotA revealed that ATRX and H3K9me3S10p are co-enriched on 25% of genomes in non-IFN treated neurons and 59% of IFNα treated neurons during Phase I (Fig 5E). These data suggest ATRX genomes are still capable of becoming phosphorylated even when bound by ATRX, but ATRX association is not lost upon following a histone phospho/methyl switch during Phase I reactivation. Together, our data indicate that ATRX functions as a restriction factor against HSV-1 reactivation by remaining bound to viral genomes even following the initial epigenetic changes known to occur during Phase I of reactivation (a phospho/methyl switch) and prevents Phase I lytic gene expression. Further, ATRX and H3K9me3 association increase when neurons are pre-treated with type I IFN to mediate the innate immune memory response in neurons that restricts HSV-1 reactivation.

## Discussion

Based on analysis of bulk cultures of ganglia, it was known that different histone post-translation modifications including H3K9me2, H3K27me3, and H3K9me3, are enriched on the latent viral genome (Cliffe et al., 2009; Kwiatkowski et al., 2009; Nicoll et al., 2016; Wang et al., 2005). However, how histone post-translational modifications arise and whether individual viral genomes with different histone PTMs contribute to the differential abilities of viral genomes to reactivate was unknown. The restriction of latency is affected by neuronal states such as stress and immune signaling (Dochnal et al., 2023; Suzich et al., 2021), but whether these processes impacted the epigenetics of the viral genome was unclear. Here, we demonstrate that immune signaling at the time of infection impacts viral heterochromatin by promoting more H3K9me3-association and restricting reactivation. Furthermore, we demonstrate mechanistically how this form of heterochromatin is refractory to reactivation via the epigenetic reader ATRX. These data are important because they show that a certain type of epigenetic structure on a latent viral genome is more deeply silenced, and this deeper form of silencing can be induced via initial infection in the presence of type I IFN.

Prior to this study, there was little information describing the nature of histone readers that are maintained at latent HSV-1 genomes. The H3K9me3 readers include the HP1 family proteins (Bannister et al., 2001; Jacobs et al., 2001; Lachner et al., 2001), ATRX (alpha-thalassemia/mental retardation, X-linked) (Dhayalan et al., 2011; Noh et al., 2015), CHD4 (Mansfield et al., 2011), TRIM66 (Jain et al., 2020), PHRF1 (Jain et al., 2020), UHRF1 (Nady et al., 2011), TNRC18 (Zhao et al., 2023), MPP8 (Chang et al., 2011), and Tip60 (Sun et al., 2009). Our data suggest that ATRX is an important H3K9me3 reader associated with the viral genome. Further work will be required to determine the extent to which other H3K9me3 readers associate with the HSV-1 genome and their impact on latency and reactivation. As we showed, ATRX is highly abundant in neurons and displays a differential localization in comparison to non-neuronal cells. Our observations suggest ATRX remains bound to regions of heterochromatin following histone phosphorylation and confirms a previous study indicating that ATRX is essential to protect the genome of neurons in times of cell stress (Noh et al., 2015). We can now extend this role for ATRX and show that it also protects neurons from viral reactivation. We also demonstrate that ATRX maintains heterochromatin silencing on the HSV-1 genome independently of DAXX, as DAXX depletion at latent time points had no impact on reactivation. This contrasts with lytic infection of HSV-1, where ATRX and DAXX function together (Lukashchuk & Everett, 2010). Here we demonstrate that the enrichment and mechanism of ATRX during HSV-1 infection in neurons is distinct. This distinct role for ATRX may help explain why HSV-1 latency is exclusively established in neurons. While our findings demonstrate a role for ATRX in maintaining HSV-1 latency, further research is needed to determine whether ATRX also contributes to the establishment of silencing during HSV-1 neuronal infection and, if so, whether it acts independently of DAXX and PML.

PML-NBs are typically abundant in non-neuronal cells but are much less abundant in the mouse nervous system, with *Pml* mRNA and protein levels significantly downregulated in post-mitotic neurons (Gray et al., 2004) (Regad et al., 2009). Previously, showed that PML-NBs were undetectable in post-natal and adult primary neurons cultured in vitro from the superior cervical and trigeminal ganglia (Suzich et al., 2021). These findings are consistent with reports of regional variability and subpopulations of neurons lacking PML signals (Catez et al., 2012) (Hall et al., 2016). Here, we describe that oxygen conditions also impact the formation of PML-NBs. We find that PML-NBs are only formed for a short period after IFNα stimulation and do not persist, nor colocalize, with the latent viral genome during latency when neurons are cultured *in vitro* in biologically relevant oxygen conditions. Other studies have observed at latency *in vivo* (28 days post-infection), the HSV-1 genome and PML were co-localized with DAXX (Catez et al., 2012). Differences in prior studies may reflect variable exposure to signaling molecules like interferons. Regardless, our pared-down latency model has allowed us to parse out that PML-NBs can modulate the nature of the viral epigenetic structure during latency establishment and that ATRX can restrict HSV-1 reactivation in genomes that were once, but no longer, associated with PML-NBs.

Our findings indicate viral genomes enriched with H3K9me3 and ATRX are prevented from entering Phase I of HSV-1 reactivation. The signaling cascade that initiates Phase I of reactivation results in a JNK-dependent methyl/phospho switch (Cliffe et al., 2015). During Phase I, a burst of lytic gene expression occurs independently of histone modification removal (Cliffe et al., 2015) (Cuddy et al., 2020) (Dochnal et al., 2022) (Whitford et al., 2022). Previous work hypothesized that the H3K9me3S10p methyl/phospho switch evicted repressive readers and allowed for transcription without the removal of repressive methylation marks (Cliffe et al., 2015). In this study, we now observe that the methyl/phospho switch occurs on genomes during Phase I of reactivation but propose that ATRX remains bound and inhibits transcription. In wells treated with interferon, we demonstrate that the population of genomes associated with ATRX and H3K9me3 increases, decreasing the population of reactivation-competent genomes. Yet, even in the interferon condition, we still observe low levels of reactivation, albeit lower than in the untreated neurons. In mock conditions, we observe a limited number of genomes independently enriched with either ATRX or H3K9me3. In these independently enriched populations, it is possible that an alternative H3K9me3 reader, such as HP1, is displaced via a methyl/phospho switch, facilitating the transition to Phase I. In both the mock and interferon treatments, the subset of latent genomes lacking both ATRX and H3K9me3 may represent the reactivation-competent population. For example, it is possible for H3K27me3, a mark observed on latent viral genomes (Cliffe et al., 2009; Kwiatkowski et al., 2009), to go through a methyl/phospho switch (H3K27me3S28p) (Gehani et al., 2010), although this mark has yet to be identified on the reactivating HSV-1 genome. Additional research is required to identify the marks and readers involved in reactivation.

Full reactivation using a PI3K inhibitor or forskolin in primary neuronal models of reactivation (Cliffe et al., 2015; Cuddy et al., 2020; Dochnal et al., 2022; Hu et al., 2022; Kim et al., 2012) as well as *ex vivo* models (Whitford et al., 2022) requires the activities of lysine 9 mono- and di-methylation, and lysine 27 mono-, di-, and tri-methylation removal by histone demethylases. KDM4 (also known as JMJD2 or JHDM3), a histone demethylase with a preference for H3K9me3, also promotes HSV-1 gene expression during reactivation induced by axotomy (Liang et al., 2013). Here, using a PI3K inhibitor, we found that inhibiting KDM4 did not affect Phase II of reactivation *in vitro*. Only when ATRX was depleted did inhibiting H3K9me3 removal contribute to reactivation. This further supports the hypothesis that ATRX functions as a reader of H3K9me3 on latent genomes, preventing H3K9me3-associated genomes from reactivating. The variation in KDM4 inhibition during reactivation triggered by different stimuli suggests that the kinetics of reactivation may vary depending on the type of stimulus. Previously, differential kinetics have already been observed with axotomy, with rapid recruitment of transcription factors (Kristie et al., 1999; Whitlow & Kristie, 2009), earlier occurrence of Phase I (Whitford et al., 2022), and reactivation occurring independently of the viral transactivator, VP16 (Sears et al., 1991; Steiner et al., 1990). This implies that distinct stimuli could influence the timing, dynamics, viral and host (restriction) factors involved in reactivation. Understanding these differences could provide insights into how specific factors modulate viral latency and reactivation, potentially leading to new therapeutic strategies.

To our knowledge, this is the first report of long-lasting epigenetic changes in neurons in response to an immune stimulus. Immune memory may serve as a mechanism through which a persistent virus modulates latency and its potential to reactivate. By altering the chromatin structure of the viral genome, immune memory can suppress reactivation under certain conditions, effectively limiting viral spread. However, this same mechanism might also enable the virus to remain latent while evading immune detection, ensuring its long-term survival within the host, highlighting the balance of viral-host fitness. Memory of previous immune stimulation in a long-lived and non-replenishing cell type, could also impact the regulation of host genes, potentially modulating neural function over time. Therefore, more work is needed to determine the implications for immune memory and long-term inflammation in the nervous system. Persistent epigenetic changes driven by immune responses, such as those induced by type I interferon, could underlie the development of neuropathologies, including neurodegenerative diseases or chronic pain syndromes, highlighting the potential implications for neuronal immune memory in human health.

## Materials and Methods

### Reagents

The reagents used in this study are described in the table below (Table S1):

### Preparation of HSV-1 virus stocks

Stayput-GFP virus was made and titered using gH-complementing F6 cells as described previously (Dochnal et al., 2022). The Vero F6 cells were cultured in Dulbecco’s modified Eagle’s medium (Gibco), supplemented with 10% FetalPlex (Gemini Bio-Products) and 250 μg/mL of G418/Geneticin (Gibco).

### Primary Neuronal Cultures

Sympathetic neurons from the Superior Cervical Ganglia (SCG) of post-natal day 0–2 (P0-P2) CD1 Mice (Charles River Laboratories) were dissected as previously described (Cliffe et al., 2015). Rodent handling and husbandry were carried out under animal protocols approved by the Animal Care and Use Committee of the University of Virginia (UVA). Ganglia were briefly kept in Leibovitz’s L-15 media with 2.05 mM l-glutamine before dissociation in collagenase type IV (1 mg/ml) followed by trypsin (2.5 mg/ml) for 20 min each at 37°C. Dissociated ganglia were triturated, and approximately 10,000 neurons per well were plated onto rat tail collagen in a 24 well plate. Sympathetic neurons were maintained in CM1 (Neurobasal^®^ Medium supplemented with PRIME-XV IS21 Neuronal Supplement (Irvine Scientific), 50 ng/ml Mouse NGF 2.5S, 2 mM l-Glutamine, and Primocin). Aphidicolin (3.3 µg/ml) was added to the CM1 for the first five days post-dissection to select against proliferating cells.

### Establishment and reactivation of latent HSV-1 infection in primary neurons

Latent HSV-1 infection was established in P6-8 sympathetic neurons from the superior cervical ganglia (SCG). Neurons were cultured for at least two days without antimitotic agents prior to infection. Where indicated, neurons were pre-treated for 18 hours and then infected in the presence of IFNα (600 IU/ml) as previously described (Suzich et al., 2021). The cultures were infected with Stayput Us11-GFP at an of MOI 5 PFU/cell, (assuming 10,000 cells per well) in Dulbecco’s Phosphate Buffered Saline (DPBS) + CaCl_2_ + MgCl_2_ supplemented with 1% fetal bovine serum, 4.5 g/L glucose, and 10 μM acyclovir (ACV) for 4 h at 37°C. The inoculum was replaced with CM1 containing 50 μM ACV. 8 days post-infection, ACV was washed out and replaced with CM1 alone. Reactivation was initiated 10 days post infection by adding Reactivation media [BrainPhys with 10% fetal bovine serum, Mouse NGF 2.5S (50 ng/ml)] and 20 μM LY294002 (Tocris). Reactivation was quantified by counting GFP-positive neurons 48 hours after the reactivation trigger was added.

### Culturing and differentiation of HD10.6 cells

The human dorsal root sensory ganglion (HD10.6) cell line was a gift from the Triezenberg Lab and cultured as previously described (Thellman et al., 2017). HD10.6 cells were incubated at 37°C and 5% CO_2_ and passaged in proliferation media (advanced DMEM/F12 supplemented with glutaMAX, B-27 supplement, prostaglandin, and bFGF). Proliferating HD10.6 cells were grown on Nunc flasks coated with human fibronectin. HD10.6 cells destined for differentiation were plated onto German Glass coverslips (Cat. #72290, Electron Microscopy Science) coated with PLO and human fibronectin at a concentration of 50,000 cells/ well. 18 hours after plating, maturation media [Neurobasal^®^ Medium supplemented with PRIME-XV IS21 Neuronal Supplement, NGF, l-Glutamine, Primocin, ciliary neurotrophic factor (CNTF), glial cell-derived neurotrophic factor (GDNF), and neurotrophin-3 (NT-3)] containing doxycycline was added to induce differentiation. Following the addition of doxycycline, half-volume medium changes were performed every 3 days until cells were fixed 10 days post differentiation for immunofluorescence.

### Analysis of mRNA expression by reverse transcription-quantitative PCR (RT-qPCR)

To assess relative expression of HSV-1 lytic mRNA, total RNA was extracted from approximately 1.0 × 10^4^ neurons using the Quick RNA™ Miniprep Kit (Zymo Research) with an on-column DNase I digestion. Following extraction, mRNA underwent a reverse transcription reaction using Maxima First Strand cDNA Synthesis Kit for RT-qPCR (Fisher Scientific) and random hexamers for first-strand synthesis to produce cDNA. Equal amounts of RNA (20–30 ng/reaction) were used per cDNA reaction. qPCR was carried out using *Power* SYBR™ Green PCR Master Mix (Applied Biosystems). The relative mRNA was determined using the comparative *C*_T_ (ΔΔ*C*_T_) method normalized to mRNA levels in control samples. Viral mRNAs were normalized to mouse reference gene 18S. All samples were run in duplicate on an Applied Biosystems™ QuantStudio™ 6 Flex Real-Time PCR System and the mean fold change compared to the reference gene calculated. Primers used are described in Table S2.

### Immunofluorescence

Cells were fixed in a solution containing 1.8% methanol-free formaldehyde in CSK buffer (10 mM HEPES, 100 mM NaCl, 300 mM Sucrose, 3 mM MgCl2, 5 mM EGTA) with 0.5% Triton X-100 and 1% phenylmethylsulfonyl fluoride (PMSF) for 10 minutes. Following fixation, cells were washed three times with PBS and were then blocked for one hour using 3% bovine serum albumin. Cells were then incubated for one hour with primary antibody at room temperature with concentrations specified in Table X. Following the primary antibody treatment, the neurons were incubated with Alexa Fluor® 488-, 555-, and 647-conjugated secondary antibodies for one hour. To visualize the nuclei, cells were incubated in Hoechst stain for 30 minutes. After all incubation times, cells were washed three times with PBS.

Images were captured using a Zeiss LSM900 microscope with Airyscan capabilities at 60x. Images were processed using the ZenBlue software and then further analyzed and processed using ImageJ.

### Click Chemistry

SCGs were infected with EdC/EdA virus at an MOI of 7. Labeled virus was prepared using a previously described method (McFarlane et al., 2019). Click chemistry was carried out as described previously (Suzich et al., 2021). Following fixation protocol described for immunofluorescence, samples were incubated in samples from Click-iT EdU Alexa Flour 555 Imaging Kit (ThermoFisher Scientific, <u>C10638</u>) following the manufacturers instructions using AFDye 555 Azide Plus. Samples were incubated in this solution for thirty minutes then washed x3 in PBS. Following the click reaction, the immunofluorescence protocol was performed.

### NucSpotA Analysis

NucSpotA is part of the Mitogenie suite and was used as previously described (Francois et al., 2023). Rotation control images were generated from original channel combination images using FIJI, prior to analysis with NucSpotA. The stated co-localization thresholds were blindly calibrated by eye.

### Preparation of lentiviral vectors

Lentiviruses expressing shRNA against PML (TRCN0000314605-validated previously(Suzich et al., 2021)), DAXX (DAXX-1 TRCN0000218672, DAXX-2 TRCN0000225715), ATRX (ATRX-1 TRCN0000302074, ATRX-2 TRCN0000081910), or a control lentivirus shRNA (pLKO.1 vector expressing a non-targeting shRNA control) were prepared by co-transfection with psPAX2 and pCMV-VSV-G(Stewart et al., 2003) using the 293LTV packaging cell line (Cell Biolabs). Supernatant was harvested at 40- and 64-h post-transfection and filtered using a 45 μM PES filter. Sympathetic neurons were transduced overnight in neuronal media containing 8μg/ml protamine sulfate and 50 μM ACV.

### CUT&Tag

CUT&Tag was carried out on latent neuronal cultures (approximately 1.2 x10^5^ neurons per reaction) using the Epicypher CUTANA CUT&Tag Kit and workflow (14-1102). Antibodies used for CUT&RUN are included in Table S3. Dual-indexed DNA libraries were prepared using the same kit. Pair-ended, partial lane sequencing and de-multiplexing were carried out using NovaSeq (Novogene). Data analysis was performed using Galaxy, command line and R code, and workflow, adapted from the cited <u>tutorial</u>. The Rivanna high-performance computing environment (UVA Research Computing) was used for the command line data processing. Data was aligned to mouse (mm39) and viral [SC16; KX946970 (Rastrojo et al., 2017)] genomes using Bowtie2. Cutadapt was used to trim i5 and i7 adapters. Bam files were converted to Bed files using samtools. Bed files were converted to Bedgraph files using bedtools. Bedgraph files were normalized to total mapped reads for host (mouse), virus (SC16), and the K Met Stat panel. Bedgraph files were visualized in integrative genome viewer (IGV) viewer, exported as SVG files, and made into figures using Inkscape. CUT&Tag experiments were repeated twice for a total of 2 biological replicates. The sum of enrichment at viral promoters and genes was used using bedtools map. To compare mock-treated and interferon-treated samples, each interferon-treated replicate was normalized by dividing it by its corresponding mock-treated replicate. Promoters and genes showing a two-fold or greater increase or decrease in the enrichment sum were considered consistent. In Galaxy, Bam files were visualized using DeepTools bamcoverage with a bin size of 1 to generate bigwig files. Bigwig files were normalized to total mapped reads for host (mouse), virus (SC16), and the K Met Stat panel. Bigwig files were used for multiBigwigSummary to generate heatmaps. Spearman correlation analysis was performed using deeptools plotCorrelation on multiBigwigSummary.

### Statistical analysis

Power analysis was used to determine the appropriate sample sizes for statistical analysis. All statistical analysis was performed using Prism V10. The normality of the data was determined with the Kolmogorov-Smirnov test. Specific analyses are included in the figure legends.

