## Supplemental figure 1-5, supplemental table 1-3 for "Interferon Dependent Immune Memory during HSV-1 Neuronal Latency Results in Increased H3K9me3 and Restriction of Reactivation by ATRX"

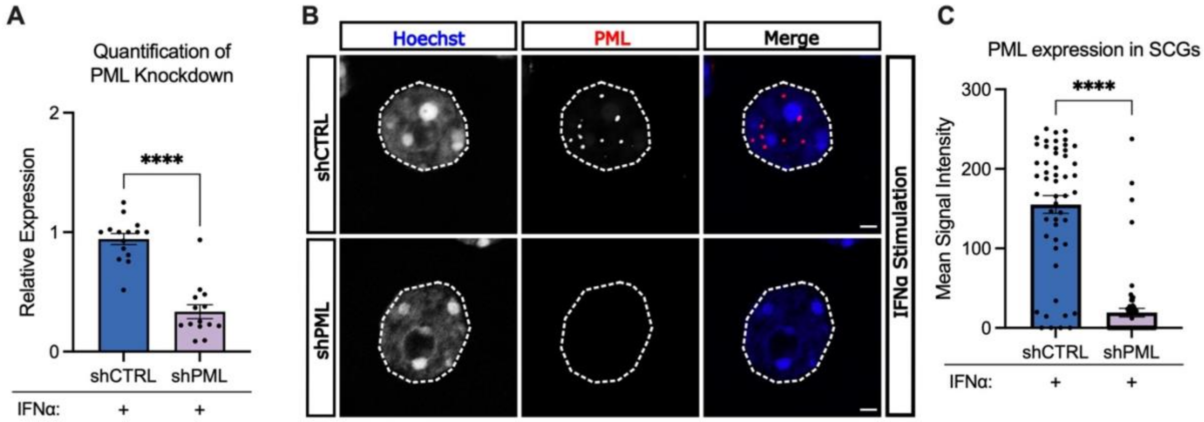


Supplemental Figure 1

**Supplemental Figure 1: Depletion of PML in neurons.**

**A)** Neurons were depleted of PML using lentivirus-mediated shRNA depletion five days prior to treatment with 600 IU/ml IFNα. Five days after knockdown, RNA was collected and *Pml* transcripts were quantified via RT-qPCR. Statistical comparisons were made using a Paired t-test. N=14 biological replicates from 5 independent dissections

**B)** Representative images of sympathetic neurons treated with 600 IU/ml of IFNα cultured in 5% oxygen. Neurons were depleted of PML and fixed 5 days later. Immunofluorescence was carried out for PML. Scale bar, 10 µm.

**C)** FIJI was used to quantify the signal intensity of PML in the nucleus 5 days post knockdown. Each data point represents one nucleus. Statistical comparisons were made using a Mann-Whitney test. N>50 biological replicates from 2 independent dissections.

Data represent the mean ± SEM. (ns not significant, *≤ 0.05, **≤ 0.01, ***≤ 0.001, ****≤ 0.0001).

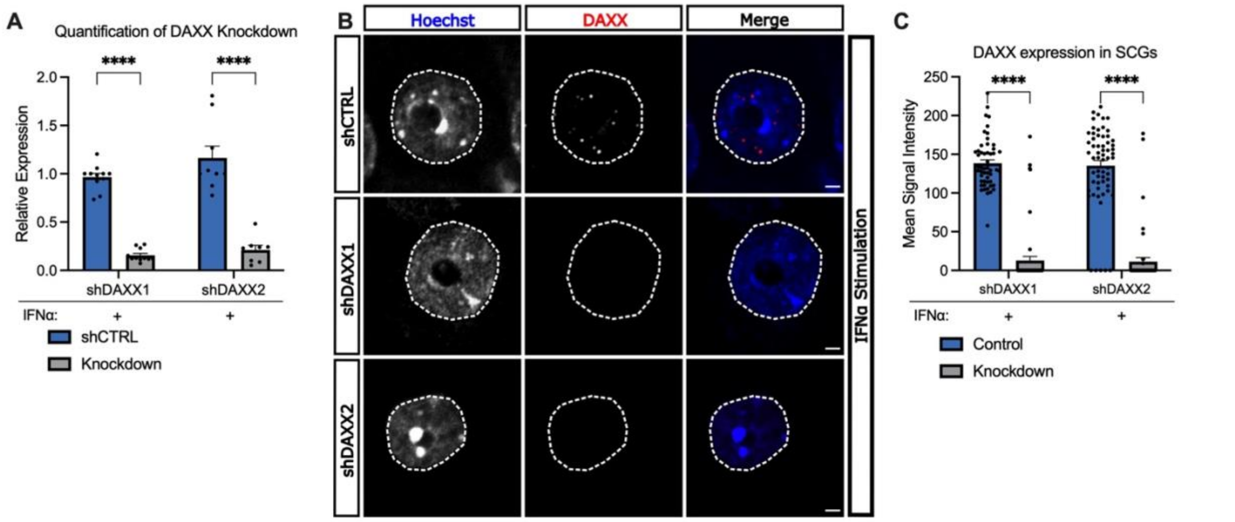


Supplemental Figure 2

**Supplemental Figure 2: Depletion of DAXX in neurons.**

**A)** Neurons were depleted of DAXX using lentivirus-mediated shRNA depletion five days prior to treatment with 600 IU/ml IFNα using two independent shRNAs. Five days after knockdown, RNA was collected, and *DAXX* transcripts were quantified via RT-qPCR. Statistical comparisons were made using a Mann-Whitney test. N>9 biological replicates from 3 independent dissections.

**B)** Representative images of sympathetic neurons treated with 600 IU/ml of IFNα cultured in 5% oxygen. Neurons were depleted of DAXX and fixed 5 days later. Immunofluorescence was carried out for DAXX. Scale bar, 10 μm.

**C)** FIJI was used to quantify the signal intensity of DAXX in the nucleus 5 days post knockdown. Each data point represents one nucleus. Statistical comparisons were made using a Mann-Whitney test. N>50 biological replicates from 2 independent dissections.

Data represent the mean ± SEM. (ns not significant, *≤ 0.05, **≤ 0.01, ***≤ 0.001, ****≤ 0.0001).

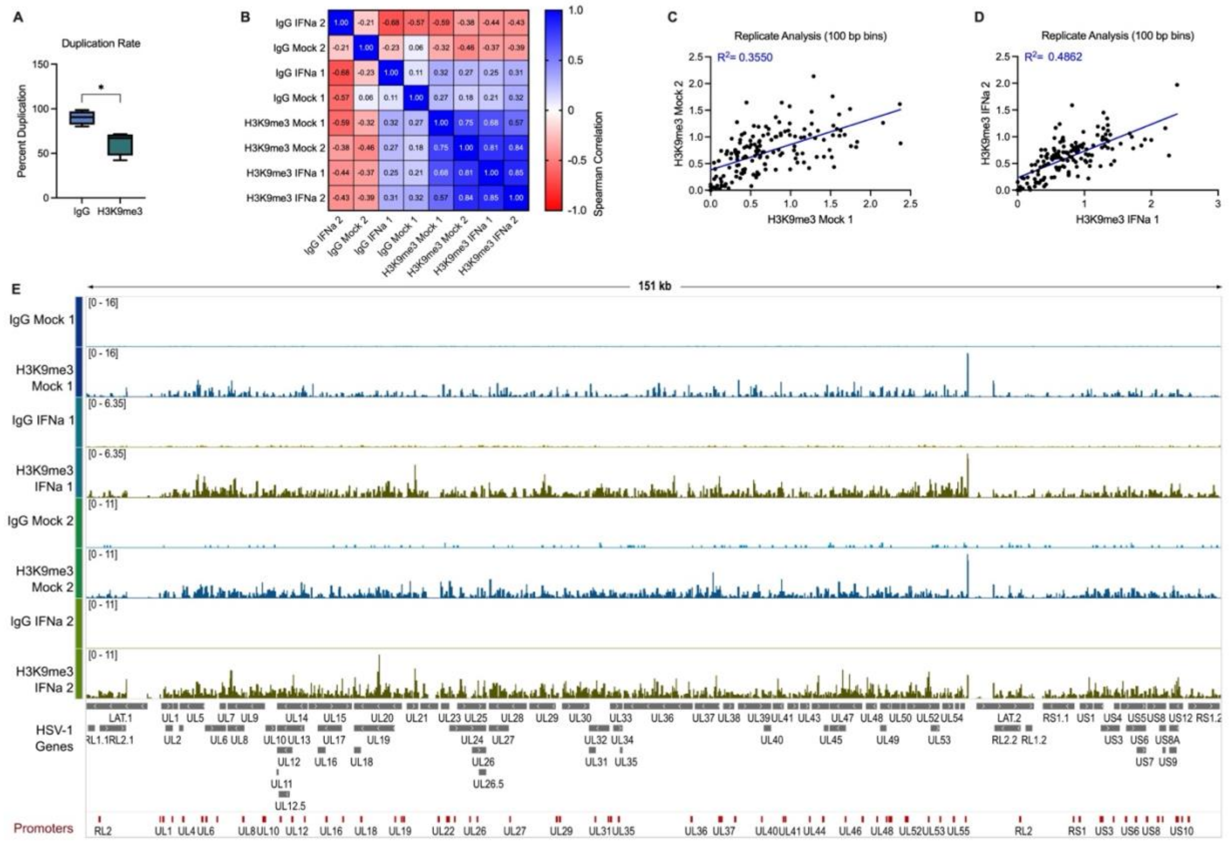


Supplemental Figure 3

**Supplemental Figure 3**

**A)** Duplication rates for viral transcripts, determined using Picard, plotted for IgG and H3K9me3.

**B, C, & D)** Normalized viral aligned bigwig files were assessed using MultiBigwigSummary in 1000 bp bins. Values within bins were used for Spearman correlation analysis (B) or plotted to analyze biological replicates in mock (C) or interferon-treated (D) conditions by linear regression analysis.

**E)** Representative integrative genome viewer images of the complete HSV-1 genome. All data was normalized to total mapped reads. Viral genes are colored gray, and viral promoters are colored red. y-axes were scaled to be group auto-scaled between replicate and treatment.

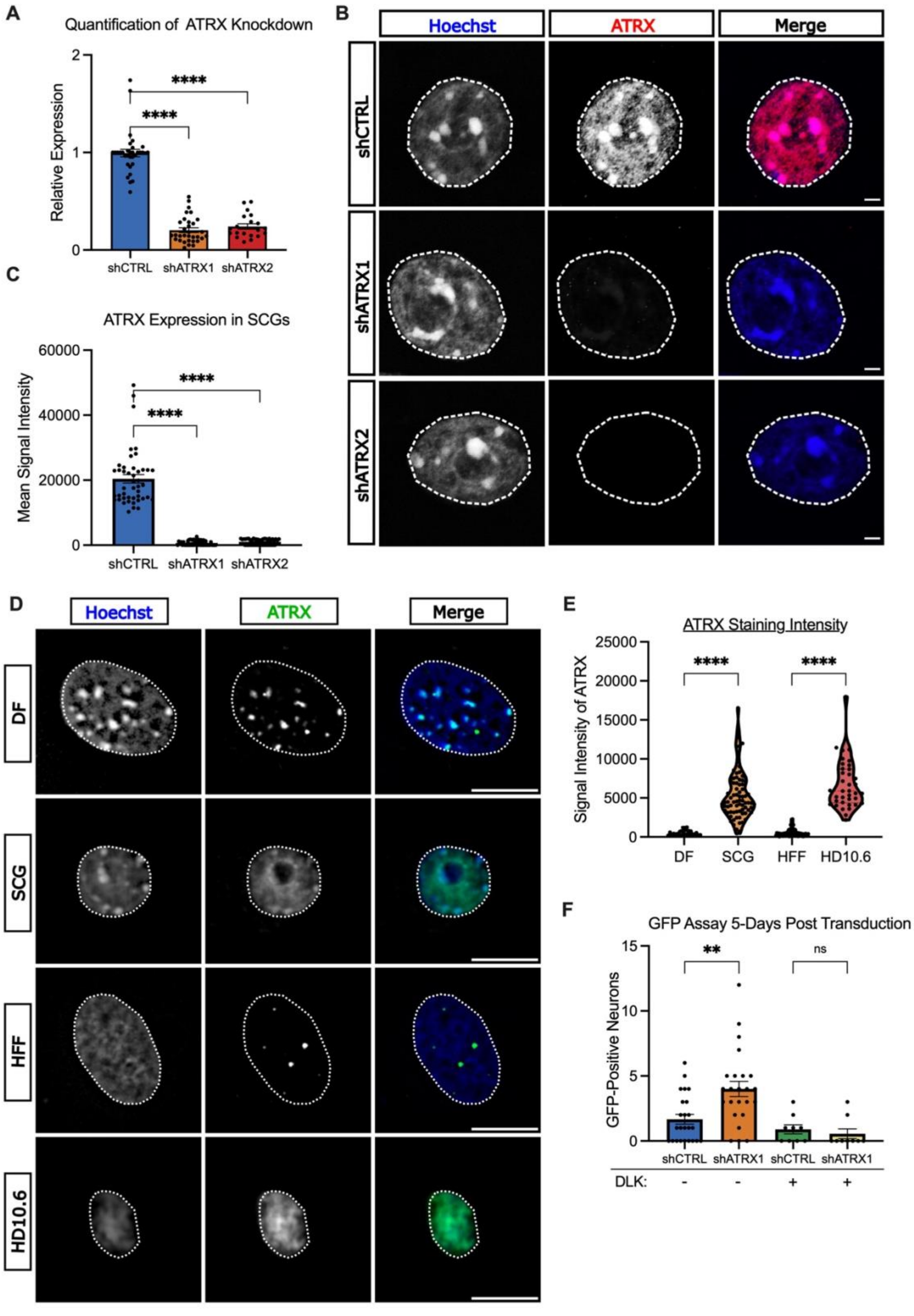


1

Supplemental Figure 4

**Supplemental Figure 4:**

**A)** Neurons were depleted of ATRX using lentivirus-mediated shRNA depletion five days prior to treatment with 600 IU/ml IFNα using two independent shRNAs. Five days after knockdown, RNA was collected, and *ATRX* transcripts were quantified via RT-qPCR. Statistical comparisons were made using an Ordinary one-way ANOVA with Tukey's multiple comparison. N>20 biological replicates from >7 independent dissections.

**B)** Representative images of sympathetic neurons treated with 600 IU/ml of IFNα cultured in 5% oxygen. Neurons were depleted of ATRX and fixed 5 days later. Immunofluorescence was carried out for ATRX. Scale bar, 10 μm.

**C)** FIJI was used to quantify the signal intensity of ATRX in the nucleus 5 days post knockdown. Each data point represents one nucleus. Statistical comparisons were made using a t-test. N>50 biological replicates from 2 independent dissections.

**D)** Representative images of primary dermal fibroblasts (DF), primary superior cervical ganglia (SCG), human foreskin fibroblasts (HFF), and human-induced peripheral neurons (HD10.6). Immunofluorescence was carried out for ATRX. Scale bar, 10 μm.

**E)** FIJI was used to quantify the signal intensity of ATRX in the nucleus of DFs, SCGs, HFFs, or HD10.6 cells. Each data point represents one nucleus. Statistical comparisons were made using a One-way ANOVA with Šídák's multiple comparisons test. N>50 biological replicates from 2 independent experiments.

**F)** Sympathetic neurons were infected with Stayput HSV-1 in the absence of IFNα and depleted of ATRX using shRNAs at 5 days post-infection. Reactivation was quantified based on the numbers Us11-GFP expressing neurons following addition LY294002 and DMSO or GNE-115 (DLK inhibitor). N=9 biological replicates from 3 independent dissections. Statistical comparisons were made using a repeated Mann-Whitney.

Data represent the mean ± SEM. (ns not significant, *≤ 0.05, **≤ 0.01, ***≤ 0.001, ****≤ 0.0001).

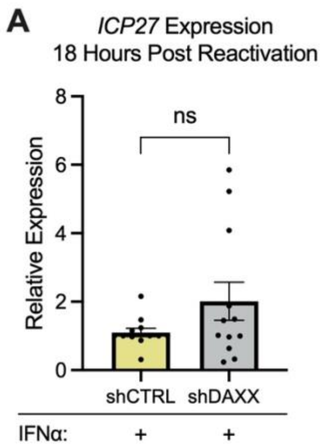


Supplemental Figure 5

**Supplemental Figure 5: DAXX does not inhibit Phase I of reactivation.**
**A)** Sympathetic neurons were infected with Stayput HSV-1 in the presence of IFNα and depleted of DAXX using shRNAs at 5 days post-infection. RNA was collected 18 hours following the addition LY294002 (Phase I). RT-qPCR was used to quantify IE (ICP27) viral gene expression. Statistical comparisons were made with a t-test. N=9 biological replicates from 3 independent dissections.

Data represent the mean ± SEM. (ns not significant, *≤ 0.05, **≤ 0.01, ***≤ 0.001, ****≤ 0.0001)

Table S1: Reagents

| Compound | Supplier | Identifier | Concentration |
|---|---|---|---|
| Aphidicolin | AG Scientific | A-1026 | 3.3 µg/ml |
| Acycloguanosine | Millipore Sigma | A4669 | 10 µM, 50 µM |
| L-Glutamic Acid | Millipore Sigma | G5638 | 3.7 µg/ml |
| LY 294002 | Tocris | 1130 | 20 µM |
| IFNα | EMD Millipore | IF009 | 600 IU/ml |
| NGF 2.5S | Alomone Labs | N-100 | 50 ng/ml |
| Primocin | Invivogen | ant-pm-1 | 100 µg/ml |
| AFDye 555 Azide Plus | Click Chemistry Tools | 1479-1 | 10 µM |
| ML-324 | Axon medchem | Axon 2081 | 10 µM |
| GDNF | Peprotrech | 450-44 | 50 ng/mL |
| PRIME-XV IS21 Neuronal Supplement | Irvine Scientific | 91142 | 1x |
| Neurobasal Medium™, minus phenol red | Gibco | 12348017 | |
| Brain Phys | STEMCELL Technologies | 05791 | |
| Advanced DMEM/F12 | Gibco-Life Tech | 12634-010 | |
| GlutaMAX (100x) | Gibco-Lif Tech | 35050061 | 1x |
| Prostaglandin E1 | Sigma | P5515 | 10ng/ml |
| fibroblast growth factor-basic (bFGF) | Stemgent | 03-0002 | 0.5ng/ml |
| Human Fibronectin | Millipore | FC010 | 5 µg/ml |
| Poly-L-ornithine (PLO) | Sigma | P4957 | 50 µg/ml |
| CNTF | Alomone | N-100 | 25ng/ml |
| GDNF | PeproTech | 450-13 | 25ng/ml |
| NT-3 | Affymetrix eBioscience | 14-8506 | 25ng/ml |
| Doxycycline | Sigma Aldrich | D9891-1G | 1ug/mL |

Table S2: Primers used for RT-qPCR

| Primer | Sequence 5'-3' |
|---|---|
| 18s F | CAC GGA CAG GAT TGA CAG ATT |
| 18s R | GCC AGA GTC TCG TTC GTT ATC |
| ICP27 F | GCA TCC TTC GTG TTT GTC ATT CTG |
| ICP27 R | GCA TCT TCT CTC CGA CCC CG |
| ICP8 F | GGA GGT GCA CCG CAT ACC |
| ICP8 R | GGC TAA AAT CCG GCA TGA AC |
| gC F | GAG TTT GTC TGG TTC GAG GAC |
| gC R | ACG GTA GAG ACT GTG GTG AA |
| ATRX F | AAA GGA AAG GGT GGG TCA TC |
| ATRX R | CTC TGT CTG CTC TGC TTC TTT |
| DAXX F | TGA CCC AGA CTC CTC GTA TTT |
| DAXX R | GTA CGG AAT TCG CTG CTC TAT G |
| PML F | GGG AAA CAG AGG AGC GAG TT |
| PML R | AAG GCC TTG AGG GAA TTG GG |

Table S3: Antibodies used and Concentrations

| Antibody | Description | Supplier | Identifier/ RRID | Concentration (IF) | Concentration (CUT&Tag) |
|---|---|---|---|---|---|
| Anti-ATRX | Rabbit polyclonal | Abcam | ab97508 | 1:100 | |
| Anti-ATRX | Mouse monoclonal | Santa Cruz Bio | sc-15408 | 1:250 | |
| Anti-Daxx | Mouse monoclonal | Santa Cruz Bio | sc-8043 / AB_627405 | 1:200 | |
| Anti-IFNAR1 | Mouse monoclonal | Leinco Tech | I-1188 / AB_2830518 | 1:1,000 | |
| Anti-H3K9me3 S10p | Rabbit polyclonal | Abcam | ab5819 | 1:200 | |
| Anti-H3K9me3 | Mouse monoclonal | Diagenode | C15200146 | 1:500 | |
| Anti-H3K9me3 | Rabbit polyclonal | Diagenode | C15410193 | 1:500 | 1:100 |
| Anti-IgG | Rabbit polyclonal | CUTANA | 13-0042 | | 1:50 |
| Anti-Murine PML | Mouse monoclonal | EMD Millipore | MAB3738 / AB_2166836 | 1:100 | |
| F(ab')2 Anti-Mouse IgG Alexa Fluor® 647 | Goat polyclonal | Thermo Fisher | A21237 | 1:1,000 | |
| F(ab')2 Anti-Mouse IgG Alexa Fluor® 555 | Goat polyclonal | Thermo Fisher | A21425 / AB_2535846 | 1:1,000 | |
| F(ab')2 Anti-Rabbit IgG Alexa Fluor® 488 | Goat polyclonal | Thermo Fisher | A11070 / AB_2534114 | 1:1,000 | |
| Anti-Chicken IgY Alexa Fluor® 647 | Goat polyclonal | Abcam | ab150175 / AB_2732800 | 1:1,000 | |
| Hoechst | | Thermo Fisher | 62249 | 1:10,000 | |
